# Prefrontal network engagement by deep brain stimulation in limbic hubs

**DOI:** 10.1101/2022.09.08.507155

**Authors:** Anusha Allawala, Kelly R Bijanki, Denise Oswalt, Raissa K Mathura, Joshua Adkinson, Victoria Pirtle, Ben Shofty, Meghan Robinson, Matthew T Harrison, Sanjay J Mathew, Wayne K Goodman, Nader Pouratian, Sameer A Sheth, David A Borton

## Abstract

**Background:** Prefrontal circuits in the human brain play an important role in cognitive and affective processing. Neuromodulation therapies delivered to certain key hubs within these circuits are being used with increasing frequency to treat a host of neuropsychiatric disorders. However, the detailed neurophysiological effects of stimulation to these hubs are largely unknown.

**Methods:** Here, we performed intracranial recordings across prefrontal networks while delivering electrical stimulation to two well-established white matter hubs involved in cognitive regulation and depression: the subcallosal cingulate (SCC) and ventral capsule/ventral striatum (VC/VS).

**Results:** We demonstrate a shared frontotemporal circuit consisting of the ventromedial PFC, amygdala, and lateral orbitofrontal cortex where gamma oscillations are differentially modulated by stimulation target. Additionally, we found subject-specific responses to stimulation in the dorsal anterior cingulate cortex and demonstrate the capacity for further tuning of neural activity using current-steered stimulation.

**Conclusions:** Our findings indicate a potential neurophysiological mechanism for the dissociable therapeutic effects seen across the SCC and VC/VS DBS targets for psychiatric neuromodulation and our results lay the groundwork for personalized, network-guided neurostimulation therapy.

## Introduction

The ability to regulate complex emotions and make controlled decisions are central to the human experience and critical for successful navigation through challenging life circumstances. Neuroimaging and electrodiagnostic studies have implicated prefrontal networks encompassing the dorsal anterior cingulate cortex (dACC), orbitofrontal cortex (OFC), ventromedial prefrontal cortex (vmPFC) and amygdala in affective and emotional regulation^1–5^, decision making and impulsivity^2,6,7^, reward evaluation^8–11^, and emotional processing^12,13^. Disruption of neural activity in these circuits is thought to lead to psychiatric disorders of mood, anxiety, and impulsivity, among other behavioral manifestations^4,14–19^. Neuromodulatory interventions^20,21^ are often used to treat such disorders, but little is known about the human *electrophysiology* of these prefrontal regions and how chronic neurostimulation therapies modify circuit dynamics underlying psychiatric symptoms. Characterizing the specific spatiotemporal prefrontal network activity implicated in affective and cognitive processing in response to therapeutic stimulation can inform stimulation paradigms on a chronic or adaptive basis and aid the prediction of an individual’s response to stimulation.

Two well-characterized affective hubs previously demonstrated to be gateways to parsimoniously engage prefrontal and corticolimbic networks through invasive means^21–24^ are the ventral capsule/ventral striatum (VC/VS) and subcallosal cingulate (SCC). The VC/VS and SCC are thought to be hubs^25^ at the crossroads of white matter pathways hypothesized to influence executive function^23^, reward processing^26,27^ and mood processing^21,28^ through their connections of varying degrees to prefrontal and limbic structures (spanning the amygdala, PFC and ACC)^29,30^ with partial overlap^31–33^. Modulation of the two targets have shown promising results in DBS studies showing improvement in symptoms of anxiety^34^, depression^21,35,36^, treatment-refractory anorexia nervosa^37^, addiction^38–40^ and obsessive compulsive disorder^41,42^. In the clinical treatment of treatment-resistant depression (TRD) with DBS, both SCC and VC/VS targets have been extensively used with mixed results. While both targets can have an antidepressant effect, responses to stimulation across the two targets are phenotypically different, and notable qualitative differences in behavioral responses (“activating” vs “calming”)^21,28,43–45^ have been observed. Prefrontal targets appear to be key in driving a response from both DBS targets^46–48^.

The aim of our study was to evaluate the network-level effects of stimulation *between* the SCC and VC/VS, two well-established targets used for psychiatric DBS therapy. We took advantage of a unique opportunity afforded through an ongoing clinical trial of DBS for TRD (NCT03437928) where we performed stimulation experiments using segmented DBS leads in the SCC and VC/VS with concurrent high-density intracranial recordings providing high spatiotemporal resolution of neural activity in two participants with TRD. We aimed to characterize the *differences* in neural response across prefrontal networks between the two DBS targets within and across patients with TRD. Given the phenotypic differences that are observable following stimulation of the VC/VS and SCC, and the role of the vmPFC, OFC, dACC and amygdala in neuropsychiatric disorders^17,49–52^, we hypothesized that we would find differentiable responses to stimulation between the DBS targets in the prefrontal cortex and that this would be unique to anatomical regions in high frequency activity (defined as 13-100 Hz for this study) or low frequency activity (defined as 1-13 Hz for our study) based on previous work implicating frequency-specific neural oscillations in mood^9,20,52,53^. Our results lay the groundwork for a more mechanistic understanding of the effects of DBS across prefrontal circuits in psychiatric disease, and better equip us to implement optimized, network-guided neuromodulation in the future.

## Methods and Materials

### Participant and study overview

Data for this study was collected from two subjects (37 year old Latino male and a 57 year old Caucasian female) diagnosed with TRD. The subjects were enrolled in an ongoing clinical trial (NCT 03437928) for DBS for TRD. Each participant gave fully informed consent according to study sponsor guidelines, and all procedures were approved by the local institutional review board at Baylor College of Medicine IRB (H-43036) prior to participation. Subjects underwent stereotactic implantation of four DBS leads (Boston Scientific Cartesia, Marlborough, MA) and 10 temporary stereo-electroencephalography (sEEG) electrodes (PMT, Chanhassen, MN) based on pre-operative scans including patient-specific tractography. Post-implantation, patients underwent a ten-day intracranial monitoring period for evaluation of brain networks involved in depression. Following the intracranial monitoring period, sEEG electrodes were removed and the four DBS leads were internalized and connected to two implanted pulse generators (IPG) (Boston Scientific Gevia, Marlborough, MA). Additional surgical details have been described previously^45^.

### Electrode implantation

Both participants were implanted bilaterally with segmented DBS leads in the VC/VS and the SCC, capable of current steering. The DBS leads used in our study consist of eight stimulation contacts: solid ring contacts at the deepest and shallow positions, as well as three-way segmented contacts located between the ring contacts. Seven total contact configurations of interest were identified per lead, including three stacked configurations listed as follows: (1) anterior-facing contacts 2 and 5, (2) posterior-left facing contacts 4 and 7 and (3) posterior-right facing contacts 3 and 6. The remaining four configurations tested were ring configurations listed as follows: (1) solid ring contact 1 (2) solid ring contact 8 (3) combination of segmented ring contacts 2, 3 and 4, and (4) combination of segmented ring contacts 5, 6 and 7 (Fig. 1b).

**Figure 1:**
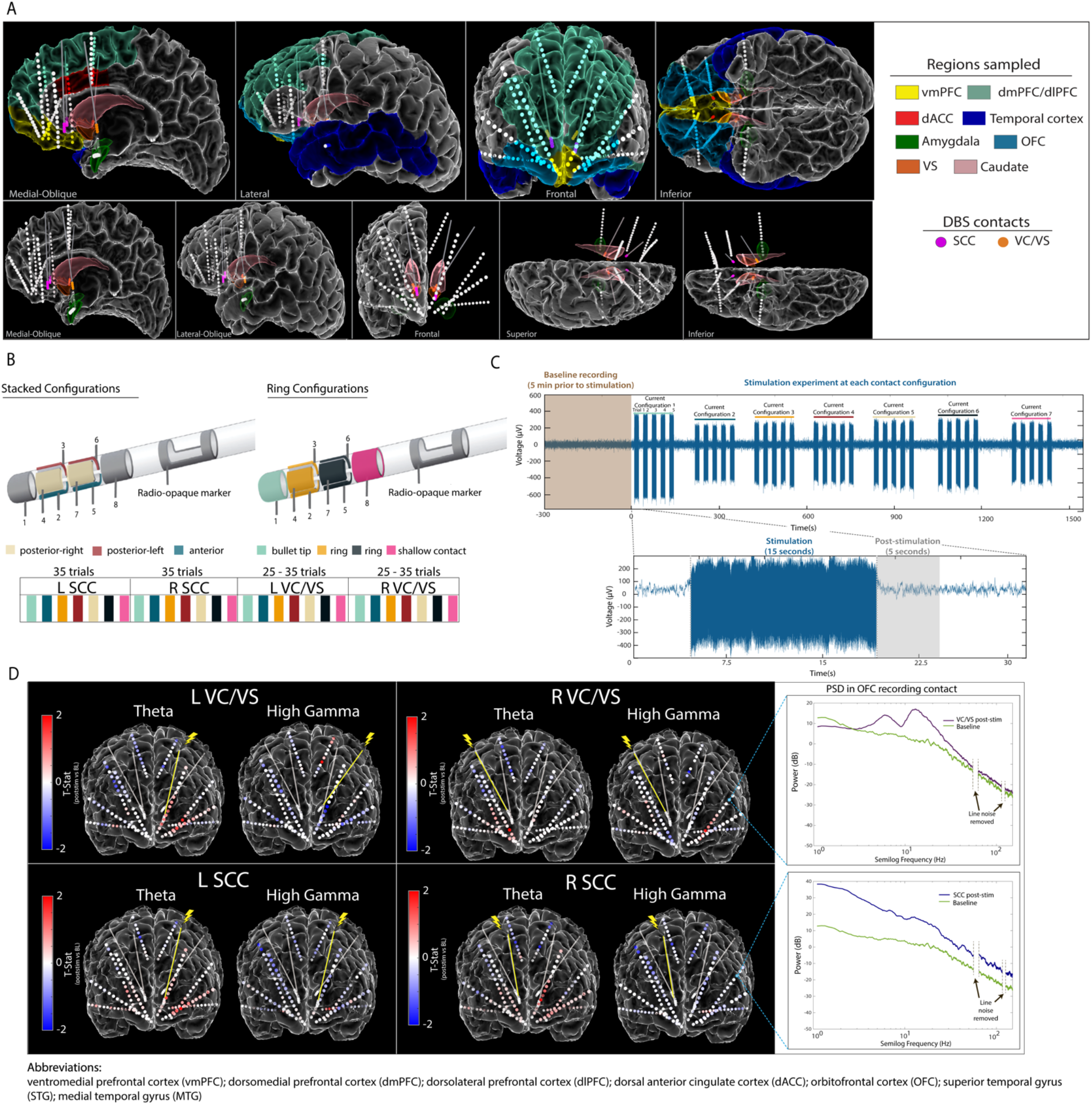
Experimental Approach. (A) Anatomical reconstruction showing placement of sEEG electrodes (top panel) and DBS leads (bottom panel). Colors in the legend (right) correspond to the region where electrodes were implanted. (B) Steerable DBS leads were used to deliver unilateral stimulation in the VC/VS and SCC respectively. Seven current configurations of interest were identified and tested across subjects. (C) Raw voltage signal recorded on an example sEEG contact during stimulation in the left SCC DBS lead. Subjects were systematically tested at each current configuration for fifteen seconds, with five trials per current configuration at each respective DBS lead. A five-second window following stimulation was used for subsequent analyses.(D) Electrode diagram showing the mean power change in theta band and high gamma band following stimulation for subject A. Each contact is colored based on the t-statistic value computed between baseline and post-stimulation for each DBS lead (red indicates increase in power following stimulation and blue indicates a decrease in power following stimulation). Inset (top-right) shows log-transformed power spectra from a recording electrode in the OFC during baseline and post-stimulation.

### Electrode Stimulation and Recording

Monopolar cathodic stimulation was delivered through each DBS lead via a Blackrock CereStim R96 (Blackrock Microsystems, Salt Lake City, UT). A stimulation amplitude of 4.8 - 5 mA was delivered at the solid ring contacts, whereas for stacked or ring configurations this amplitude was split evenly among contacts to enable current steering, never exceeding 5 mA in total or at any time a charge density of 30 µC/cm^2^. Stimulation was applied at 130 Hz, with a pulse width of 180 uS and interphase gap of 100 uS. In subject A, we tested the seven identified stimulation combinations (3 stacked configurations, 4 ring configurations) for each DBS lead in the SCC, and five combinations (3 stacked configurations, 2 ring configurations) in each DBS lead in the VC/VS. In subject B, we tested all seven combinations across each of the four DBS leads. Each trial of stimulation consisted of 15 seconds of stimulation on followed by 10 seconds without stimulation (Fig. 1c). Trials were repeated 5 times per contact configuration per DBS lead seriatim, resulting in 25-35 trials per DBS lead for subject A and 35 trials per DBS lead for subject B.

### Data Acquisition and Signal Processing

Electrophysiological signals from implanted sEEG electrode contacts were recorded using a 256-channel NeuroPort Acquisition System (Blackrock Microsystems, Utah) at a sampling rate of 2 kHz, with a hardware high pass filter applied at 0.3 Hz. Recordings from sEEG contacts were analyzed offline using custom scripts written in MATLAB (Mathworks Inc. Natick, MA) and Python. LFP signals were demeaned, decimated to 1 kHz and bandpass filtered between 1-250 Hz. A butterworth notch filter was applied to remove line noise at 60, 120 and 180 Hz, respectively. Recordings were bipolar re-referenced by subtracting the activity of adjacent electrode contact pairs. Any channels with excessive noise or without a clear neural signal were removed from the analysis. See Supplementary Methods for additional details.

## Results

The goal of our study was to quantify prefrontal network responses to intracranial stimulation between two DBS targets: the SCC and VC/VS. We first evaluated the effects of stimulation for each DBS target (pre-vs. post-stim, p-values adjusted to compensate for multiple comparisons reported in Supplementary Methods and Tables S1-4; Fig. 1c) on high-density stereo-EEG (sEEG recordings) in two subjects with TRD (Fig. 1a-b; Supplementary Methods). We then compared neural responses (see Supplementary Methods) following stimulation between the two DBS targets (SCC post-stim vs. VC/VS post-stim, adjusted p-values reported in Supplementary Tables S5-6) on high frequency neural activity (beta, low gamma and high gamma band power) and low frequency activity (delta, theta and alpha band power). A representative example of the electrode coverage is shown in Fig. 1d, illustrating bilateral modulation of low frequency power (e.g., theta) and high frequency power (e.g., high gamma) across recording contacts following unilateral stimulation in Subject A. Given the previously established roles of the dACC, amygdala, OFC and vmPFC in affective and cognitive regulation in psychiatric disorders, we focused our analyses across these key four anatomical regions and describe our findings for each key region in detail below.

### vmPFC

#### Stimulation-induced response of high frequency activity

We first sought to understand the effect of stimulation on high frequency activity in the vmPFC given its broad involvement in cognitive, affective and emotional processing^2^. Here, we found consistent differences when evaluating neural responses in high frequency bands between SCC and VC/VS stimulation (Fig. 2c,f) in both subjects. **Specifically, we found that SCC consistently increased gamma power in both subjects while VC/VS decreased gamma power**. Within Subject A, left SCC stimulation elicited a significant increase in spectral power in high gamma (pre-vs post-stim, adj.p<0.01). The response to stimulation was significantly different between both DBS targets in high gamma band (adj.p<0.01) and low gamma band (adj.p<0.01) irrespective of the hemisphere of stimulation. The inverse relationship in which SCC increased high-frequency activity and VC/VS decreased high-frequency activity was also observed in beta band (adj.p<0.001) in subject A. In Subject B (Fig. 2d), we observed right VC/VS stimulation significantly decreased low gamma power (adj.p<0.05) and beta power (adj.p<0.001) from baseline. In the same subject, neural responses were significantly different between the two DBS targets as observed in low gamma (adj.p<0.05) and high gamma (adj.p<0.01, corrected; Fig. 2f).

**Figure 2:**
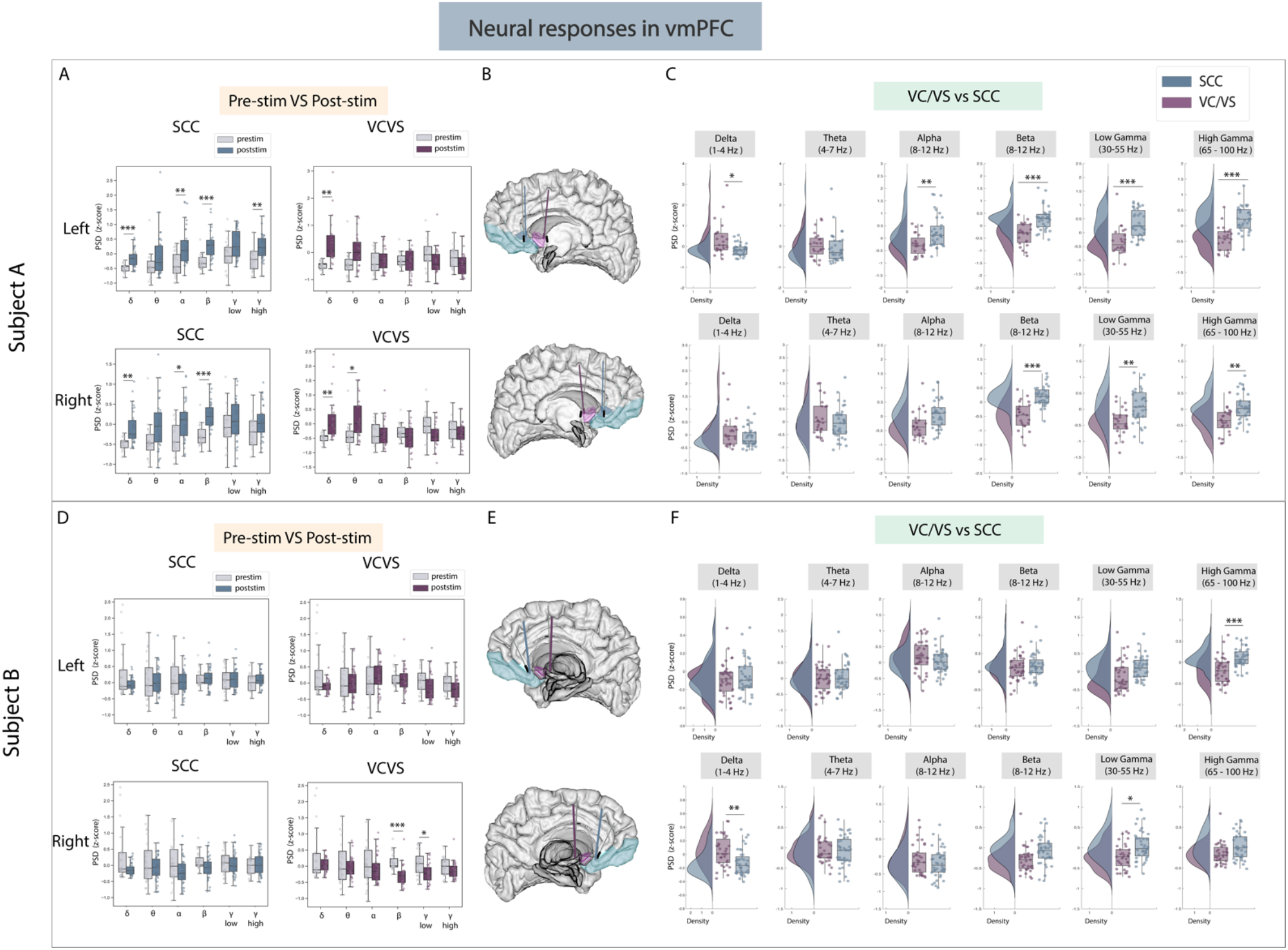
Neural responses in the vmPFC following SCC stimulation vs. VC/VS stimulation. (A) Distribution of spectral power across all post-stimulation trials vs pre-stimulation (baseline) in the vmPFC after z-scoring in six pre-defined frequency bands (delta, theta, alpha, beta, low gamma and high gamma) following SCC stimulation (left) and VC/VS stimulation (right) in subject A (B) Corresponding anatomical location of the vmPFC highlighted in light blue and corresponding VC/VS and SCC DBS leads highlighted depending on hemisphere of stimulation. Stimulation in left hemisphere is on top, while stimulation in right hemisphere is shown on the bottom. (C) Distribution of spectral power across six pre-defined frequency bands contrasting neural responses following SCC stimulation and VC/VS stimulation. (D)-(F) Replicate of figures in panels (A)-(C) for subject B. * indicates significance where adj.p-value < 0.05, corrected; ** indicates significance where adj.p-value ≤ 0.01, corrected; *** indicates significance, where adj.p-value ≤ 0.001, corrected.

#### Stimulation-induced response of low frequency activity

We next explored if the same opposing response between SCC and VC/VS was observed in low-frequency activity. We did observe a significant difference in delta power between SCC and VC/VS stimulation in both subjects (Fig. 2c,f). While both SCC and VC/VS stimulation significantly increased delta power from baseline (adj.p<0.01) respectively, we found that VC/VS stimulation drove a larger increase in delta power than SCC stimulation and the responses between the two DBS targets were significantly different (adj.p<0.05) in subject A. In subject B, it appears that while both SCC and VC/VS stimulation drive a decrease in delta power from baseline (Fig. 2d), the response between the two targets is still significantly different, where VC/VS drives a smaller decrease than SCC stimulation (adj.p<0.01, corrected; Fig. 2f).

### Amygdala

#### Stimulation-induced response of high frequency neural activity

The next area of interest for this study was the amygdala, given its role in emotional regulation^12^. In the amygdala, **while both VC/VS and SCC stimulation elicited increases in low and high gamma power, responses in both low and high gamma were still significantly different between the DBS targets in both subjects** (Fig. 3c,f). SCC stimulation significantly increased high frequency activity from baseline in both subjects (Fig. 3a,d). In subject A, SCC stimulation drove a significant increase in beta (adj.p<0.001), low gamma (adj.p<0.001) and high gamma power (adj.p<0.001). In subject B, right SCC stimulation significantly increased low gamma (adj.p<0.001) and high gamma power (adj.p<0.001) while significantly decreasing beta power (adj.p<0.01). VC/VS stimulation also significantly increased high gamma power in both subjects (adj.p<0.05, corrected; Fig 3. a,d). In subject A, right VC/VS stimulation also significantly increased low gamma power (adj.p<0.05) and in subject B, left VC/VS stimulation significantly decreased beta power (adj.p<0.05). When contrasting neural responses between the two DBS targets (Fig. 3c,f) we observed that SCC increased low gamma (adj.p<0.05) and high gamma power (adj.p<0.05) significantly higher than VC/VS stimulation, and right SCC increased beta power significantly higher than VC/VS stimulation (adj.p<0.01) in subject A. In subject B, beta power and high gamma power were again significantly higher (adj.p<0.05) following right SCC stimulation compared to right VC/VS stimulation, and low gamma power was significantly higher (adj.p<0.001) following SCC stimulation compared to VC/VS stimulation irrespective of the hemisphere of stimulation.

**Figure 3:**
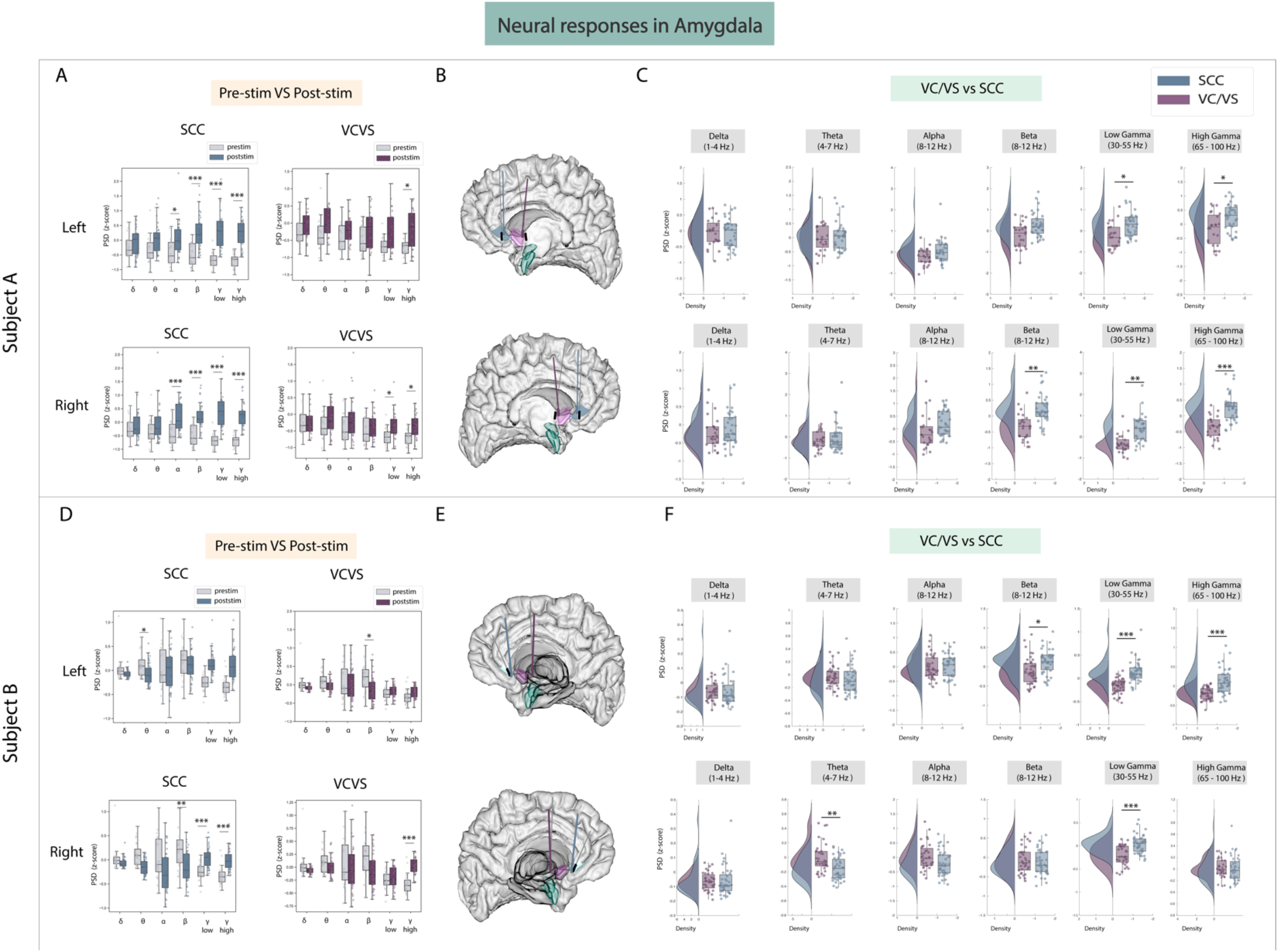
Neural responses in the amygdala following SCC stimulation vs. VC/VS stimulation. (A) Distribution of spectral power across all post-stimulation trials vs pre-stimulation (baseline) in the amygdala after z-scoring in six pre-defined frequency bands (delta, theta, alpha, beta, low gamma and high gamma) following SCC stimulation (left) and VC/VS stimulation (right) in subject A (B) Corresponding anatomical location of the amygdala highlighted in green and corresponding VC/VS and SCC DBS leads highlighted depending on hemisphere of stimulation. Stimulation in left hemisphere is on top, while stimulation in right hemisphere is shown on the bottom. (C) Distribution of spectral power across six pre-defined frequency bands contrasting neural responses following SCC stimulation and VC/VS stimulation. (D)-(F) Replicate of figures in panels (A)-(C) for subject B. * indicates significance where adj.p-value < 0.05, corrected; ** indicates significance where adj.p-value ≤ 0.01, corrected; *** indicates significance, where adj.p-value ≤ 0.001, corrected.

#### Stimulation-induced response of low frequency activity

We were similarly interested to see if significant differences between SCC and VC/VS were prevalent in low frequency activity. We found responses to SCC and VC/VS stimulation were significantly different in theta band in subject B (adj.p<0.01, corrected; Fig. 3c,f). Here, SCC drove a decrease in power relative to the VC/VS. However, no consistent modulation of low frequency activity across the two subjects was otherwise observed.

### Lateral and medial OFC

#### Stimulation-induced response of high and low frequency activity (lateral OFC)

The lOFC was a third target of interest because it has been implicated in cognitive and reward processing and recently employed as a target for neuromodulation to improve mood^44,52^. We found neural responses to SCC stimulation were significantly different from VC/VS stimulation in subject A (adj.p<0.05, corrected; Fig. 4c) and once again followed the same inverse relationship seen in the amygdala, **where SCC drove an increase in power in beta, low gamma, and high gamma bands relative to the VC/VS**. In the same subject, SCC stimulation also significantly increased beta, low gamma and high gamma power from baseline (adj.p<0.01). In subject B, the significant difference in response between SCC and VC/VS stimulation was observed in high gamma power (adj.p<0.01, corrected; Fig. 4f) following stimulation in the left hemisphere.

**Figure 4:**
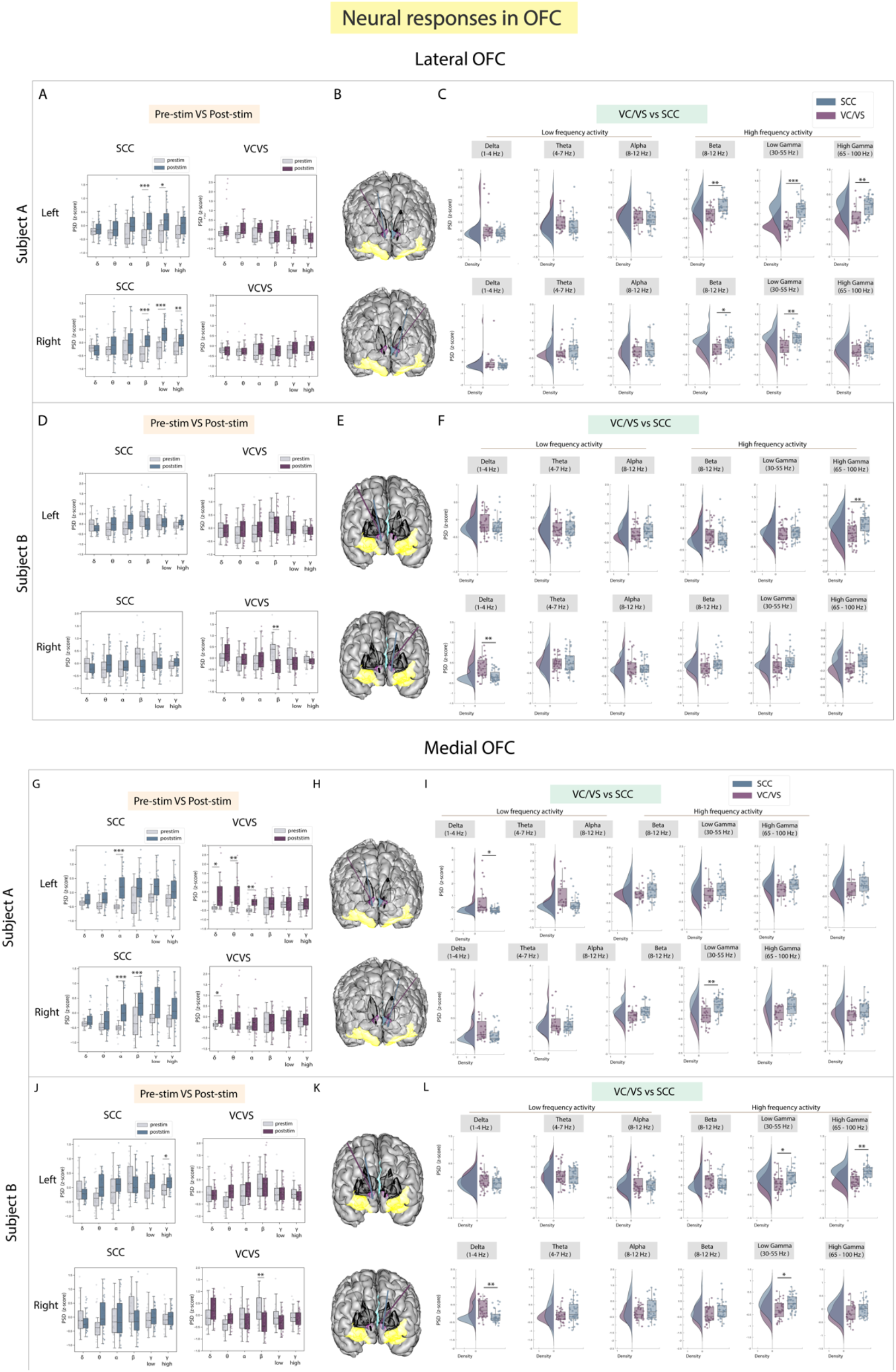
Neural responses in the OFC following SCC stimulation vs. VC/VS stimulation. (A) Distribution of spectral power across all post-stimulation trials vs pre-stimulation (baseline) in the lOFC after z-scoring in six pre-defined frequency bands (delta, theta, alpha, beta, low gamma and high gamma) following SCC stimulation (left) and VC/VS stimulation (right) in subject A (B) Corresponding anatomical location of OFC highlighted in yellow and corresponding VC/VS and SCC DBS leads highlighted depending on hemisphere of stimulation. Stimulation in left hemisphere is on top, while stimulation in right hemisphere is shown on the bottom. (C) Distribution of spectral power across six pre-defined frequency bands contrasting neural responses following SCC stimulation and VC/VS stimulation. (D)-(F) Replicate of figures in panels (A)-(C) for subject B. (G) Distribution of spectral power across all post-stimulation trials vs pre-stimulation (baseline) in the mOFC after z-scoring in six pre-defined frequency bands following SCC stimulation (left) and VC/VS stimulation (right) in subject A. (H) Corresponding anatomical location of OFC highlighted in yellow and corresponding VC/VS and SCC DBS leads highlighted depending on hemisphere of stimulation. Stimulation in left hemisphere is on top, while stimulation in right hemisphere is shown on the bottom. (I) Distribution of spectral power across six pre-defined frequency bands contrasting neural responses following SCC stimulation and VC/VS stimulation. (J)-(L) Replicate of figures in panels (G)-(I) for subject B. * indicates significance where adj.p-value < 0.05, corrected; ** indicates significance where adj.p-value ≤ 0.01, corrected; *** indicates significance, where adj.p-value ≤ 0.001, corrected.

We next explored the neural responses of low frequency activity to DBS across the two targets.

Surprisingly, we did not find many differences in neural responses to SCC and VC/VS stimulation. Delta band power was significantly modulated, (adj.p<0.01): right VC/VS stimulation increased delta power and right SCC stimulation decreased delta power. The differential delta band power change was, however, subject specific.

#### Subject-specific response to stimulation observed in high and low frequency neural activity (medial OFC)

We explored stimulation response in the mOFC separately as the lateral and medial orbitofrontal structures have shown to have distinct roles in cognitive and reward processing^7^. We found that differences between responses in high frequency activity following SCC vs. VC/VS stimulation were subject-specific in the mOFC (Fig. 4i,l). For example, the inverse relationship where SCC increases high frequency activity and VC/VS decreases high frequency activity was observed in subject A. Here, right SCC stimulation significantly increased beta power from baseline (adj.p<0.001, corrected; Fig. 4g), and we observed a significant difference in response to stimulation in beta power between the SCC and the VC/VS (adj.p<0.01, corrected; Fig. 4i). In subject B, SCC significantly increased high gamma power from baseline (adj.p<0.05, corrected; Fig. 4j). The inverse relationship between SCC and VC/VS stimulation-induced responses of high frequency activity was observed in low gamma (adj.p<0.05) and high gamma (adj.p<0.01), where SCC stimulation increased activity relative to VC/VS stimulation (Fig. 4l).

When assessing stimulation response in low frequency activity in the mOFC, we found significant differences between responses to VC/VS vs. SCC stimulation seen across both subjects in delta band (Fig. 4i,l). In subject A, left VC/VS stimulation significantly increased power in delta band, and this response was significantly higher than the neural response following SCC stimulation (adj.p<0.05). In subject B, right VC/VS stimulation significantly increased delta power relative to SCC stimulation (adj.p<0.01).

### dACC

#### Subject-specific response to stimulation observed in high and low frequency neural activity

A key region known to play an important role in cognitive control and emotional processing is the dACC^3,6^. When examining high frequency activity in the dACC, we found that significant differences following stimulation between the SCC and VC/VS were also individual-specific in the dACC but still followed the inverse relationship between the two DBS targets observed in high frequency activity in other ROIs (Fig. 5c,f). Right SCC stimulation significantly increased beta power (adj.p<0.01) compared to VC/VS stimulation in subject A. In subject B, left SCC stimulation significantly increased low gamma power from baseline (adj.p<0.01) while stimulation of either hemisphere in the SCC increased high gamma power (adj.p<0.001, corrected; Fig. 5d). Left VC/VS stimulation similarly significantly increased low gamma power (adj.p<0.05) while stimulation of either hemisphere significantly increased high gamma power from baseline (adj.p<0.01, corrected; Fig. 5d). However, the responses between left SCC and left VC/VS stimulation were still significantly different in low gamma band (adj.p<0.01;Fig 5f) and SCC stimulation drove a larger increase in low gamma power relative to the VC/VS.

**Figure 5:**
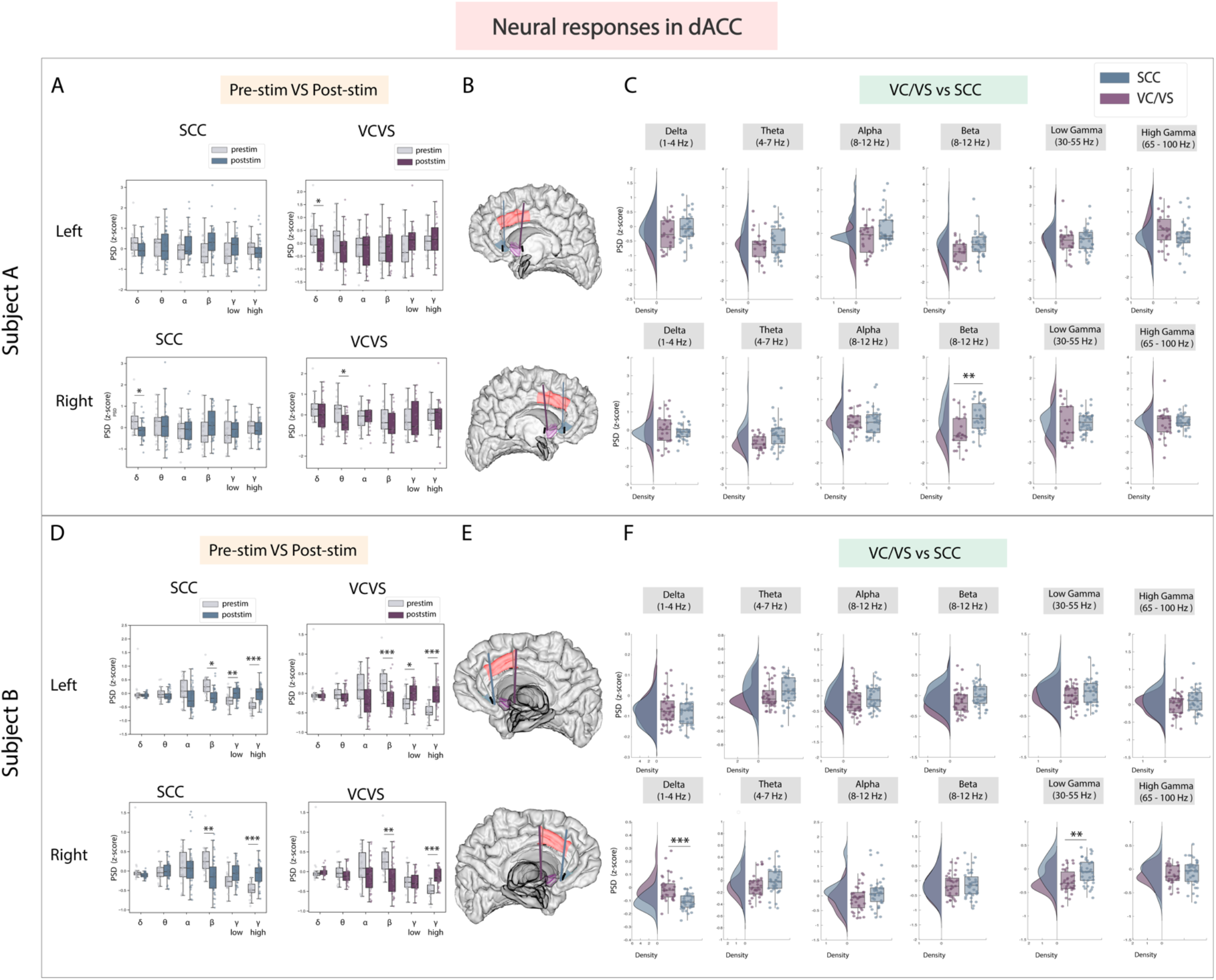
Neural responses in the dACC following SCC stimulation vs. VC/VS stimulation. (A) Distribution of spectral power across all post-stimulation trials vs pre-stimulation (baseline) in the dACC after z-scoring in six pre-defined frequency bands (delta, theta, alpha, beta, low gamma and high gamma) following SCC stimulation (left) and VC/VS stimulation (right) in subject A (B) Corresponding anatomical location of dACC highlighted in red and corresponding VC/VS and SCC DBS leads highlighted depending on hemisphere of stimulation. Stimulation in left hemisphere is on top, while stimulation in right hemisphere is shown on the bottom. (C) Distribution of spectral power across six pre-defined frequency bands comparing neural responses between SCC stimulation and VC/VS stimulation. (D)-(F) Replicate of figures in panels (A)-(C) for subject B. * indicates significance where adj.p-value < 0.05, corrected; ** indicates significance where adj.p-value ≤ 0.01, corrected; *** indicates significance, where adj.p-value ≤ 0.001, corrected.

When examining low frequency activity in response to stimulation in the dACC, we also observed a significant difference between VC/VS and SCC stimulation in delta power (adj.p<0.001) in subject B - similar to the pattern observed in other ROIs - where SCC decreased power in a low frequency band (delta) and VC/VS stimulation increased power. In subject A, right SCC stimulation significantly decreased delta power (adj.p<0.05) but no significant difference was observed between SCC and VC/VS stimulation in delta power.

#### Current steering can drive dissociable changes across prefrontal networks

To examine whether current steering enabled greater specificity in modulating neural features of interest, we evaluated spectral power for each tested contact configuration within each segmented DBS lead. Given the few trials per contact configuration for each lead, a full statistical investigation controlling for multiple comparisons was not possible. Instead, our results highlights some of the largest differences observed for a pair of DBS leads (in this case, right SCC and VC/VS) when examining the effect of current steering on neural activity to demonstrate potential effects that can be explored more fully in future work. The first feature selected was alpha-band power (see Supplementary Methods), where we found the largest amount of variance across contact configurations in the right SCC (Fig. 1b). Current-steered stimulation in right SCC differentially increased or decreased alpha power across ROIs (Fig. 6a, left). A significant effect of current steering was seen in the lOFC (p<0.05, uncorrected) and mOFC (p<0.01, uncorrected). Across the right SCC DBS lead, two stacked configurations (posterior electrodes 2-5 and 3-6) and a ring configuration (electrodes 2-3-4) demonstrated a decrease in alpha power in the OFC whereas remaining configurations resulted in an increase in power (the largest mean increase was seen in response to the bullet tip electrode, electrode 1). In response to current-steered stimulation in right VC/VS lead (Fig. 1b), mean differences were observed in the dACC and amygdala depending on the contact configuration (Fig. 6a, right), but there was no significant effect of current steering.

**Figure 6:**
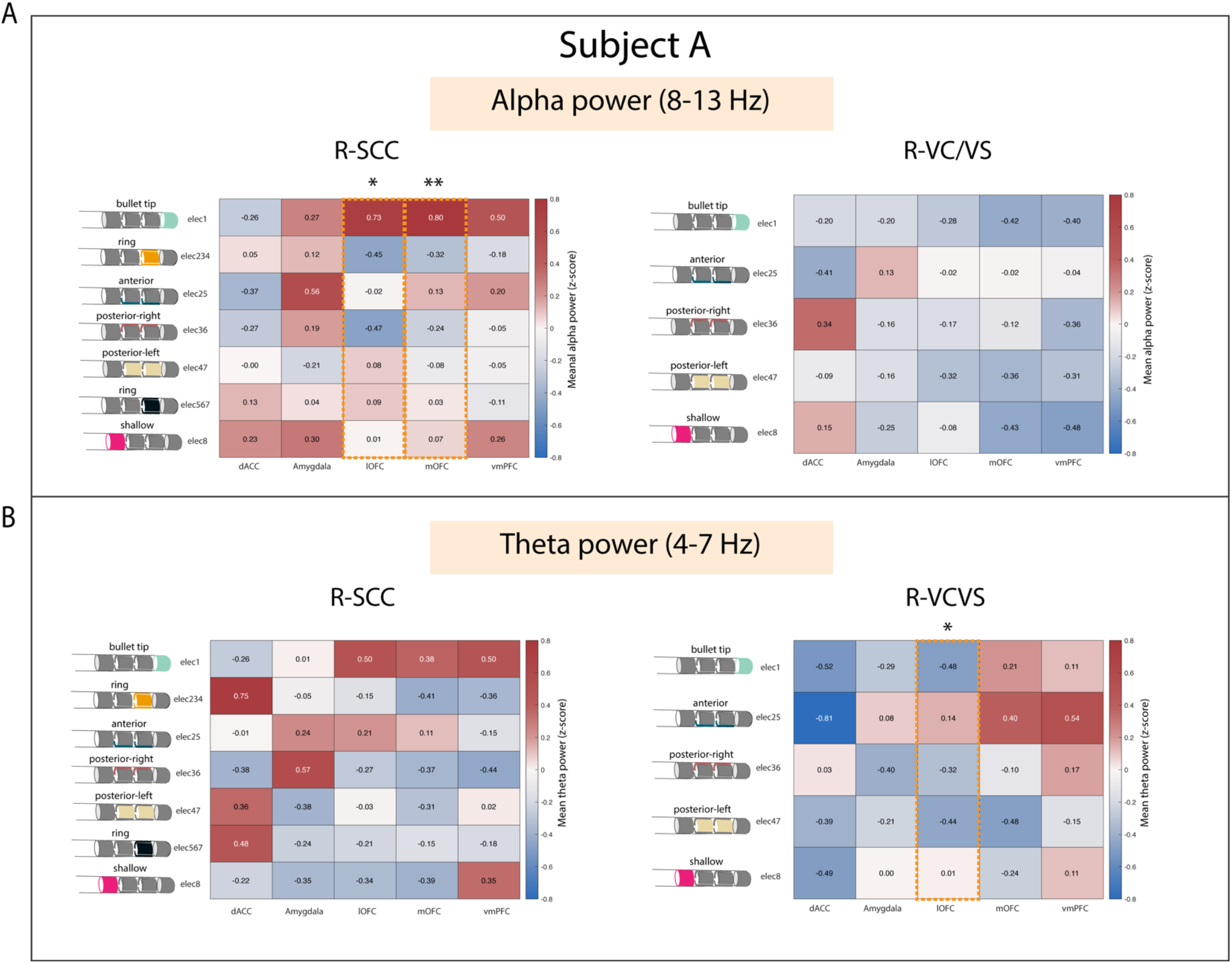
Current steering elicits differential effects in neural features implicated in depression. (A) Mean spectral power in alpha band in response to current steered stimulation in the right SCC (left) and right VC/VS (right) in subject A. Each row corresponds to a tested contact configuration across the respective DBS lead, while each column represents mean spectral power for each ROI. Seven current configurations were tested in the SCC while five were tested in the VC/VS for subject A. Alpha band was selected based on the neural feature that showed the greatest variance across all ROIs following current steered stimulation across the right SCC DBS lead. (B) Mean spectral power in theta band in response to current steered stimulation in the right SCC (left) and right VC/VS (right) in subject A. Each row corresponds to a tested contact configuration across the respective DBS lead, while each column represents mean spectral power for each ROI. Seven current configurations were tested in the SCC while five were tested in the VC/VS for subject A. Alpha band was selected based on the neural feature that showed the greatest variance across all ROIs following current steered stimulation across the right VC/VS lead.

Our second feature of interest was theta-band power, that showed the largest amount of variance across contact configurations tested in the right VC/VS DBS lead. A significant effect of current steering across the right VC/VS lead was seen only in the lOFC (p<0.05, uncorrected; Fig. 6b, right). In the lOFC, the bullet tip electrode (electrode 1) configuration and two stacked configurations (posterior electrodes 3-6 and 4-7) decreased power while the anterior stacked configuration (electrodes 2-5) increased mean power. More broadly, mean differences were observed across other ROIs such as the vmPFC and mOFC between contact configurations as shown Fig. 6b (right) following VC/VS and SCC stimulation, respectively.

## Discussion

Through a unique intracranial stimulation and recording dataset collected in two subjects with TRD, we obtained results with three main conclusions (Fig. 7). First, we demonstrate that two canonical targets for psychiatric neuromodulation, the SCC and VC/VS, elicit network-wide neurophysiological responses in both high and low frequency activity following stimulation. Second, as hypothesized, we show that stimulation in the SCC and VC/VS drive differentiable neural responses. Specifically, we observed opposite effects on gamma activity in the vmPFC, and differing degrees of modulation on gamma activity in the lOFC and amygdala, where SCC stimulation consistently drives a greater increase in gamma oscillations relative to the VC/VS. Third, we highlight preliminary evidence of subject-specific neurophysiological responses between stimulation targets, and the ability to further modify neural responses using current-steered DBS in the OFC.

**Figure 7:**
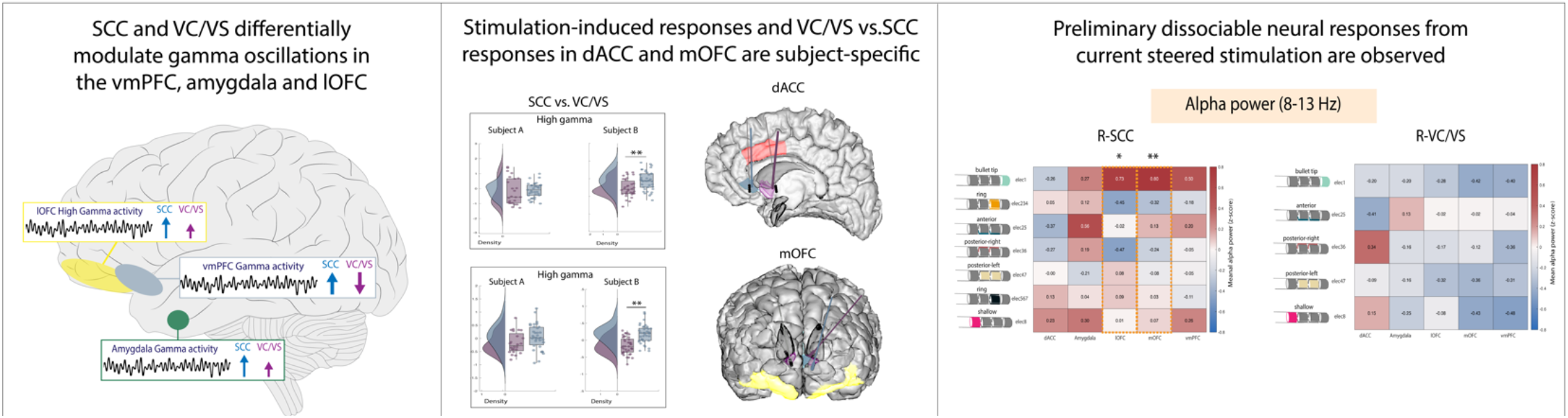
Summary of results.

Previous tractography work demonstrates that projections from the SCC and VC/VS overlap in the amygdala and medial PFC, but the anatomical trajectory and pattern of connectivity of these projections are distinct^32,33^, including the sub-regions that receive projections from the SCC and VC/VS respectively^33^. The difference in connectivity patterns may partially account for the distinct patterns of gamma activity in the vmPFC and amygdala between the two stimulation targets. The modulation of gamma activity seen in the amygdala is additionally supported by a recent study implementing amygdala gamma power as a biomarker for closed-loop VC/VS DBS in a case study for TRD^20^. Interestingly, our results also show modulation of gamma activity in the amygdala following stimulation in *both* targets, but to differing degrees.

Previous studies have also demonstrated anatomical connectivity between the SCC and the OFC, and the VC/VS and the OFC^54,55^, and initial results from our group have shown differing effective connectivity between the SCC and VC/VS to the lOFC respectively in TRD participants^56^, leading us to expect differences in neural responses between VC/VS and SCC stimulation. While we observed differing degrees of gamma power modulation in the lOFC depending on the DBS target stimulation in both subjects, we did not always observe an overlap in neural responses to stimulation between lOFC and mOFC which might be explained in part by previous work indicating the distinct roles of the lateral vs. medial OFC^18,52^.

While both VC/VS and SCC stimulation can ameliorate depressive symptoms, they have been described to modulate different dimensions of affective processing and mood: VC/VS stimulation has been reported to increase motivation and energy^28,57^, while SCC stimulation has reportedly increased calmness, alertness and exteroceptive awareness^43,58^. It is possible that the dissociable increase/decrease in gamma activity in the vmPFC and the differing degree of gamma modulation in the amygdala and lOFC may be underlying the differences in SCC vs. VC/VS stimulation described in acute behavioral reports seen elsewhere and further work will elucidate our understanding of this phenomenon.

In the dACC, we expected consistent differences in gamma power modulation following SCC and VC/VS stimulation, given recent work implicating dACC gamma power in positive affective behaviors^53^ and differential connectivity of the dACC to the VC/VS^54,59^ and SCC^55^. However, differences in gamma responses between SCC and VC/VS stimulation were subject-specific. Additionally, within a given DBS lead, we observed subject-specific responses between post-stim and pre-stim in the dACC, as well as all other ROIs. The observed subject-specific stimulation responses both between and within the VC/VS and SCC suggest that increasing efforts to personalize therapy may rely on these within-subject electrophysiological signatures across networks to deliver optimized stimulation. Efforts utilizing subject-specific biomarkers for psychiatric DBS have been recently successfully demonstrated^20^, alongside network-guided neuromodulation for psychiatric disorders^60^.

Future efforts incorporating behavioral measures corresponding to functional domains within mental illness, alongside electrophysiological measurements to build brain-behavior relationships with stimulation will help identify generalizable principles that can be potentially extended to sub-domains of MDD in the broader population^61^. **Even in the absence of behavioral measurements, characterization of stimulation-induced neural response during resting state, especially between DBS targets, enables tailoring of therapy (i.e. selecting an optimal DBS target and stimulation paradigm) for disorders such as depression, based on a patient’s neural response to stimulation, putative biomarker of mood, or another functional domain**.

Current-steered DBS provides an added parameter for differentially modulating implicated networks and neural biomarkers. A finer grained approach to precisely target anatomical regions implicated in psychopathology of depression for a desired behavioral responses is needed^62^, and the degree of stimulation-induced neural response may plausibly determine a patient’s therapeutic response. However, the topography is less well-established across psychiatric targets, in contrast to movement disorders where tracts around stimulation targets are well-understood^63^. Future study is needed to understand the extent of the effect that directionality may have on connectivity across prefrontal networks.

Primary limitations of this study include the small sample size (N=2), and lack of randomization of stimulation conditions within or across the two DBS targets. While a rigorous pipeline for optimal surgical targeting was implemented^45^, one possible reason for inconsistent results within and across subjects is that small variations in targeting may result in modifications in the electrophysiological effects observed. As we were concerned about insufficient time for stimulation washout between trials and stimulation parameters, the baseline window used for analysis was a five-minute recording collected prior to stimulation experiments. To address possible temporal autocorrelation in the baseline recording, we performed a correction procedure (Supplementary Fig. S2). However, delta power in vmPFC and lOFC in Subject B had a larger amount of autocorrelation during the baseline recording that could not be fully corrected with our approach, thus, results for those specific neural features must be viewed provisionally.

## Acknowledgments

We thank our study participants for their dedication and commitment to this study. This study was supported by the National Science Foundation Graduate Research Fellowship (ABA), National Institutes of Health grant no. F99 NS124181 (ABA), UH3 NS103549 (SAS, DAB, ABA, KRB, DO, RKM, JA, VP, SJM, WKG and NP), grant no. S10 OD025181 (DAB), grant no. K01 MH116364 (KRB), grant no. NIH R01 MH127006 (KRB), grant no. R01 274MH114854 (WKG), and the McNair Foundation (SAS)

## Author Contributions

SAS and KRB initiated the study. ABA, KRB, SAS, DAB designed the experiment. ABA conceptualized analysis procedures, analyzed the data, and drafted the manuscript with support from MH, KRB, SAS and DAB. SAS, NP, VP, WKG, SJM and KRB oversaw the organization of the clinical trial, subject recruitment, and regulatory activities. SAS and NP performed neurosurgical procedures in the subjects’ clinical trials. ABA, RKM, KRB, JA and DO performed data collection in the neurophysiology monitoring unit. RKM, BS, MR and KRB performed MRI analysis. ABA, SAS, and DAB wrote the manuscript, and all authors had an opportunity to review the manuscript and provide intellectual feedback.

## Disclosures

SAS has consulting agreements with Boston Scientific, Neuropace, Abbott, and Zimmer Biomet. NP is a consultant for Second Sight Medical Products, Abbott Laboratories, Boston Scientific, and Sensoria Therapeutics. WKG has received donated devices from Medtronic, has consulted for Biohaven Pharmaceuticals and receives royalties from Nview, LLC. SJM is supported through the use of resources and facilities at the Michael E. Debakey VA Medical Center, Houston, Texas and receives support from The Menninger Clinic. SJM has served as a consultant to Allergan, Alkermes, Axsome Therapeutics, BioXcel Therapeutics, Clexio Biosciences, COMPASS Pathways, Eleusis, Engrail Therapeutics, Greenwich Biosciences, Intra-Cellular Therapies, Janssen, Levo Therapeutics, Perception Neurosciences, Praxis Precision Medicines, Neumora, Neurocrine, Relmada Therapeutics, Sage Therapeutics, Seelos Therapeutics, and Sunovion. He has received research support from Biohaven Pharmaceuticals, Boehringer-Ingelheim, Janssen, Merck, Sage Therapeutics, and VistaGen Therapeutics.

The remaining authors declare no competing interests.

This article is currently available as a preprint on bioarxiv. Doi: https://doi.org/10.1101/2022.09.08.507155

