## Supplementary Info for "Prefrontal network engagement by deep brain stimulation in limbic hubs"

Allawala et al.

**Supplementary Figures**

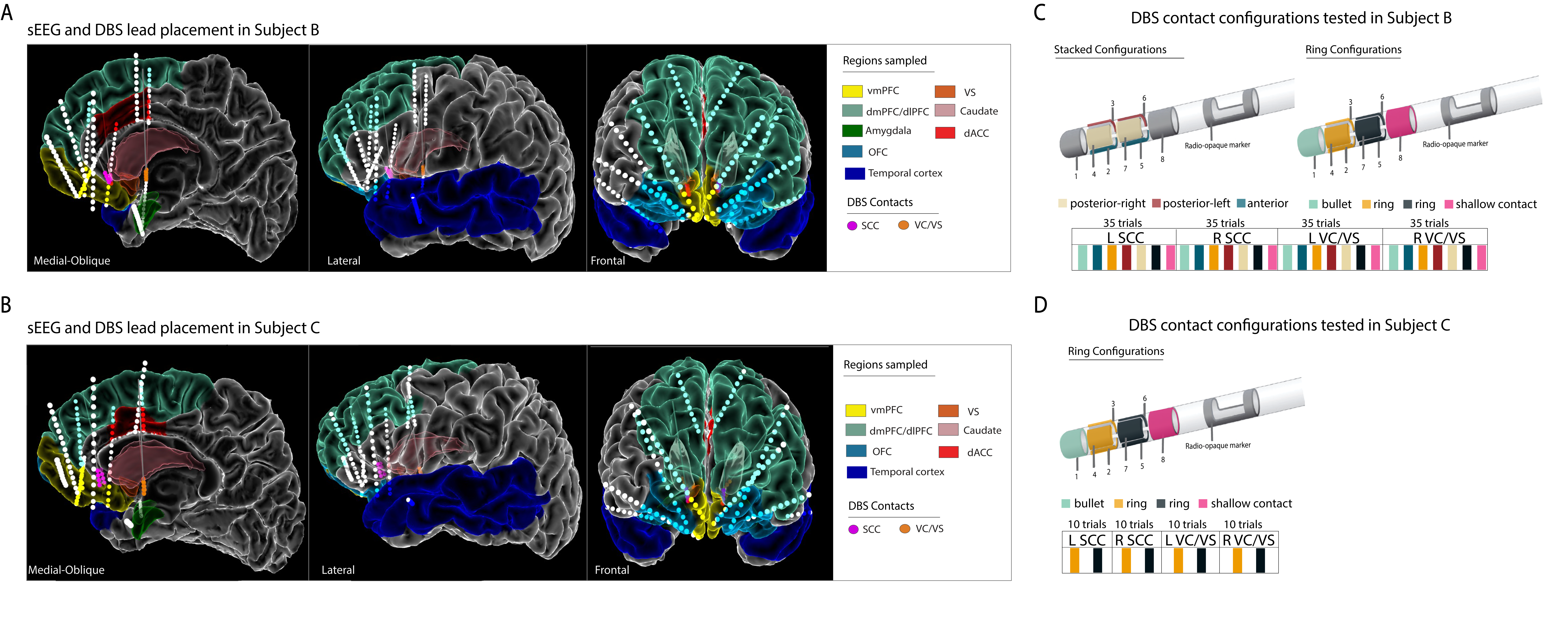

**Supplementary Figure S1:** Location of sEEG and DBS electrodes and tested stimulation configurations in Subject B.

(A - B) Anatomical locations of sEEG contacts and DBS leads within Subject B. Both subjects had recording sEEG contacts across the vmPFC, dlPFC, dACC, OFC and Temporal cortex. (C) Contact configurations tested on each DBS lead. A total of 7 current configurations were tested across each DBS lead, resulting in 35 trials of stimulation per lead.

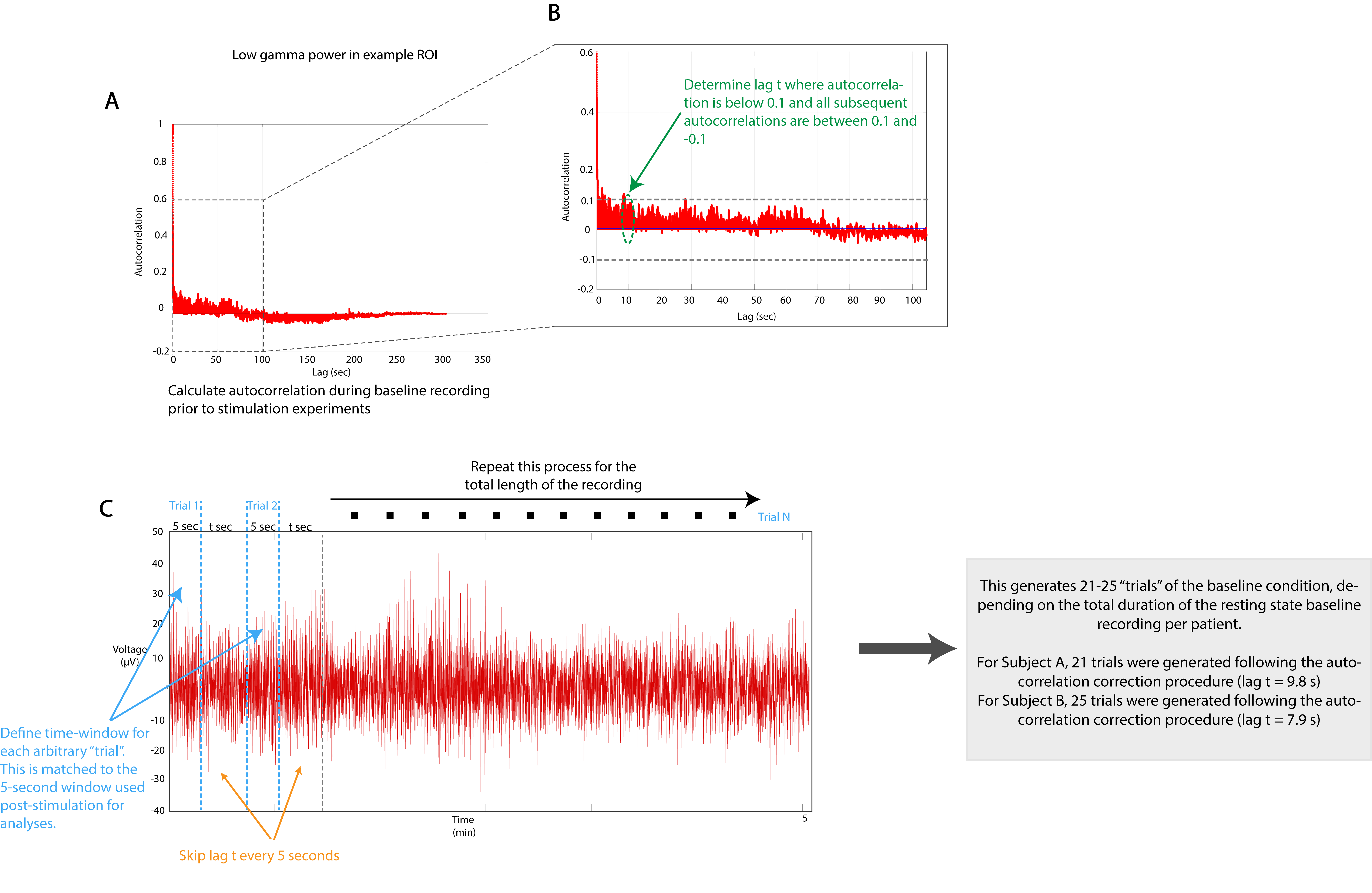
**Supplementary Figure S2:** Autocorrelation correction procedure for baseline recordings.

We performed a correction procedure to account for any autocorrelation in our baseline recordings. We took 5-second windows (to match the length of time for each trial in the post-stimulation data) and skipped t seconds between them to allow for any existing autocorrelation to disappear, and so that we could model surrogate “trials” as independent (which is necessary for permutation testing). In order to choose the appropriate lag t, we performed the following steps: we first computed power spectral estimates across the entire recording, and using the autocorr.m function in MATLAB, calculated the autocorrelation for each region of interest (ROI) and frequency band of interest within each subject. For example, autocorrelation values during the baseline recording of low gamma power in an example ROI for Subject A is shown in (A). We then identified a lag t for which the autocorrelation is below 0.1 and all subsequent autocorrelation values are below 0.1 and greater than -0.1 for all frequency bands and ROIs (indicated by dashed lines in (B)). Additionally, for each subject we ensured that lag t that was larger than the majority of the lags identified across all regions and frequency bands, ensuring that the number of samples is the same across all ROIs and neural features. We observed a few exceptions in a few neural features and ROIs where this procedure could not account for autocorrelation – in these cases the autocorrelations were extended for such long lags that we would have to discard a significant amount of data. In such cases, we still used our lag t for these exceptions and results for these exceptions can be viewed provisionally. The exceptions are listed as follows: delta band in vmPFC and lOFC in subject B. The steps outlined in A-C are performed for each subject separately.

**Supplementary Methods**

*Electrode Implantation*

Intracranial sEEG electrodes for local field potential (LFP) recordings were implanted bilaterally across several cortical and subcortical targets based on previous work implicating their roles in mood, reward, as well as cognitive and affective processing^1–4^. Regions sampled included the dorsolateral prefrontal cortex (dlPFC), ventromedial prefrontal cortex (vmPFC), dorsal anterior cingulate cortex (dACC), lateral and medial orbitofrontal cortex (lOFC, mOFC), superior frontal gyrus (SFG), superior and medial temporal gyri (STG, MTG) and the amygdala (Fig. 1a). Postoperative CT scans and pre-operative MRI scans were aligned using the Functional Magnetic Resonance Imaging for the Brain Software Library's (FMRIB's) Linear Image Registration Tool (FLIRT). Electrode coordinates were manually determined from the co-registered CT in BioImage Suite and placed into native MRI space. The reconstructed cortical surface, segmented cortical and subcortical structures and electrode coordinates were visualized using the Multi-Modal Visualization Tool^5^.

*Data Acquisition and Signal Processing*

Electrode contacts in gray matter located in the regions of interest were identified and pre-processed signals were then grouped (averaged) per region of interest. The location of electrode contacts in gray matter vs white matter and the labels for anatomical region of interest was verified by visual review of the MRI and CT by an expert rater (BS).

To evaluate the response of the sampled networks before and after stimulation, we analyzed the spectral power in the 5 seconds following stimulation to avoid artifact contamination. We identified a window of 600 ms post-stimulation that was additionally excluded from analysis to avoid residual post-stimulation artifacts in the signal. The multitaper spectral estimation method was used to extract power spectral density (PSD) from the sEEG recording using the *mspectrumc.m* function from the Chronux toolbox^6^. Spectral power was then averaged within standard frequency bands (delta = 1-4 Hz, theta = 4-8 Hz, alpha = 8-12 Hz, beta = 12-30 Hz, low-gamma = 30-55 Hz, high-gamma = 65-100 Hz). The spectral power across the baseline windows and post-stimulation windows across all stimulation experiments were z-scored within subjects.

Prior to starting the stimulation experiments, 5 minutes of baseline recording was collected for each subject (Fig. 1c). To avoid temporal autocorrelation, the autocorrelation was computed for spectral power within each frequency band of interest and region of interest, across time (Supplementary Fig. S3). The number of lags *t* where the autocorrelation was at or below 0.1 and all subsequent autocorrelation values were between 0.1 and -0.1. Lag *t* was then used to generate surrogate “trials” from the baseline recording, where *t* seconds was skipped every 5 seconds. The resulting 5-second trials were used for analysis to compare against the post-stimulation windows during the stimulation experiments (additional details in Supplementary Fig. S3).

*Statistical Analysis*

To test the difference between pre- (baseline) and post-stimulation, non-parametric permutation testing was performed on the z-scored data using custom scripts written in MATLAB. Data labels from post-stimulation and baseline windows were randomly shuffled, and then the absolute value of the t-statistic for a two-sample, pooled variance, t-test was computed for each pair of shuffled data. This procedure was repeated 1000 times. We used a single-step maxT procedure to correct for multiple tests^7,8^, namely, each absolute t-statistic was compared to the distribution (over permutations) of the maximal absolute t-statistic across all regions of interest and frequency bands of interest in order to obtain corrected p-values that control the familywise error rate (corrected p-values reported in Supplementary Tables S1-4 and uncorrected p-values are reported in Supplementary Tables S8-11).

To test the difference between PSD changes across the network following unilateral VC/VS stimulation versus SCC stimulation, non-parametric permutation testing was performed on the z-scored data using custom scripts written in MATLAB. We compared the effect of unilateral VC/VS to unilateral SCC stimulation (i.e. left VC/VS was tested against left SCC stim and right VC/VS stim was tested against right SCC stim). Data labels for unilateral SCC stimulation and unilateral VC/VS stimulation were randomly shuffled, and then the absolute value of the t-statistic for a two-sample, pooled variance, t-test was computed for each pair of shuffled data. This procedure was repeated 1000 times. We used a single-step maxT procedure to correct for multiple tests^7,8^, namely, each absolute t-statistic was compared to the distribution (over permutations) of the maximal absolute t-statistic across all regions of interest and frequency bands of interest in order to obtain corrected p-values that control the familywise error rate (corrected p-values reported in Supplementary Table S5-6 and uncorrected p-values are reported in Supplementary Tables S12-13).

The permutation testing procedure and single-step maxT procedure is described in detail as follows:

Let $M$ be a $d\times n$ matrix where $M_{ij}$ denotes the measurement of the $i$th feature (e.g., $\alpha$-power on recording electrode 3) on the $j$th trial. Let $C$ be a $1\times n$ vector where $C_{j}\in\{1,2\}$ denotes which of two conditions the $j$th trial corresponds to (e.g., pre-stim $=1$, post-stim $=2$). $M$ is held fixed throughout, so we suppress it in our notation, but all of the statistics defined below operate on $M$. We do not suppress $C$ in our notation, since we will be permuting the entries of $C$ for the permutation tests described below. Let $T(C)$ be a $d\times1$ vector where the $i$th entry $T_{i}(C)$ is a statistic for comparing trials in condition $1$ to trials in condition $2$ as indicated by $C$ on the basis of the $i$th feature measured in $M$. In the following our statistic will be the absolute value of the classical test-statistic used in the two-sample (unpaired) t-test assuming equal variances, namely,

$$T_{i}\left( C \right)=\frac{\left| \hat{\mu}_{i1}\left( C \right)-\hat{\mu}_{i2}(C) \right|}{\sqrt{\left( \frac{1}{n_{1}}+\frac{1}{n_{2}} \right)\left( \frac{{(n}_{1}-1)s_{i1}^{2}(C)+\left( n_{2}-1 \right)s_{i2}^{2}(C)}{n-2} \right)}}$$

where $n_{b}$ is the number of trials with condition $b\in\{1,2\}$, namely,

$$n_{b}=\#\{j:C_{j}=b\}$$

where $\hat{\mu}_{ib}$ is the sample mean of the measurements of the $i$th feature in condition $b\in\{1,2\}$, namely,

$$\hat{\mu}_{ib}(C)=\frac{1}{n_{b}}\sum_{j:C_{j}=b} M_{ij}$$

and where $s_{ib}^{2}$ is the sample variance of the measurements of the $i$th feature,

$$s_{ib}^{2}\left( C \right)=\frac{1}{n_{b}-1}\sum_{j:C_{j}=b} \left( M_{ij}-\hat{\mu}_{ib} \right)^{2}$$

For clarity, we note that $n_{1}+n_{2}=n$ and that both $n_{1}$ and $n_{2}$ depend on $C$, but not on the order of the entries in $C$, so we have suppressed that dependence in our notation.

If $\pi=\left( \pi_{1},\ldots,\pi_{n} \right)$ is a permutation of the numbers $(1,\ldots,n)$, then we define the $1\times n$ vector of permuted condition labels $C^{\pi}$ via $C_{j}^{\pi}=C_{\pi_{j}}$. If $\pi$ is a random permutation (chosen uniformly from all possible $n!$ permutations), then $T_{i}(C^{\pi})$ uses the above two-sample t-test statistic to compare trials randomly assigned to condition $1$ to trials randomly assigned to condition $2$ on the basis of the measurements of feature $i$. The vector $T\left( C^{\pi} \right)$ contains the results of this comparison for each of the $d$ features. Importantly, each entry of $T(C^{\pi})$ uses the *same* permutation $\pi$.

Consider the null hypothesis $H_{i}$ that the measurements of the $i$th feature are unrelated to the condition labels in $C$. (More formally, the null hypothesis is that $M_{i1},\ldots,M_{in}$ are conditionally exchangeable given $C$.) A Monte Carlo permutation test can be used to create valid p-values for each of the null hypotheses $H_{1},\ldots,H_{d}$ as follows. We sample a total of $L$ independent random permutations $\pi^{k}=(\pi_{1}^{k},\ldots,\pi_{n}^{k})$ for $k=1,\ldots,L$. We use $L=1000$ in the paper. For each such permutation we compute the t-statistic vector $T^{k}=T(C^{\pi^{k}})$. We also compute the t-statistic vector for the original $C$, namely, $T^{0}=T(C)$. A valid p-value for $H_{i}$ is

$$p_{i}=\frac{1}{L+1}\sum_{k=0}^{L} \mathbb{1}\left[ T_{i}^{k}\geq T_{i}^{0} \right]$$

where the indicator function $\mathbb{1}[A]$ returns $1$ if $A$ is true and $0$ if $A$ is false. This p-value gives the fraction of times (among all permutations and the original) that the test statistic for feature $i$ is at least as large as the original observed test statistic for feature $i$. It is important that the $k=0$ case is included in the sum and that we use $\geq$ and not $>$ in the comparison to ensure the validity of the test. Note that the smallest possible value of $p_{i}$ is $1/(L+1)$. These $p_{i}$ are the uncorrected or unadjusted p-values used in the paper.

To control for multiple hypothesis tests, we use a single-step version of the max-T procedure that will strongly control the familywise error rate (FWER) across all features subject to a minor caveat described below. For the $k$th permutation (including the case $k=0$ corresponding to no permutation), define the maximum statistic as

$$T_{*}^{k}=\max_{i=1,\ldots,d} T_{i}^{k}$$

which is a number, not a vector. A valid adjusted p-value for $H_{i}$ that controls the FWER over the collection $H_{1},\ldots,H_{d}$ is

$$p_{i}^{*}=\frac{1}{L+1}\sum_{k=0}^{L} \mathbb{1}\left[ T_{*}^{k}\geq T_{i}^{0} \right]$$

This adjusted p-value gives the fraction of times (among all permutations and the original) that the maximum test statistic over all features is at least as large as the original observed test statistic for feature $i$. We always have $p_{i}^{*}\geq p_{i}$, but equality is definitely possible (unlike, say, Bonferroni adjusted p-values). These $p_{i}^{*}$ are the corrected or adjusted p-values used in the paper. Our rationale for using max-T adjusted p-values instead of Bonferroni adjusted p-values is that max-T adjusted p-values can have much better power for positively correlated p-values (which we expect to see in our context) and also require many fewer permutations to achieve good power.

Unlike the Bonferroni correction, the max-T procedure requires a technical assumption called *subset pivotality* to ensure strong control of the FWER. Roughly speaking, the subset pivotality assumption states that the joint distribution of all features where the null hypothesis is true does not change across conditions. Consider, for instance, the following scenario. The distribution of feature $1$ is unchanged between condition $1$ and condition $2$ so that $H_{1}$ is true. Similarly, the distribution of feature $2$ is unchanged between condition $1$ and condition $2$ so that $H_{2}$ is true. But features $1$ and $2$ are more correlated in condition $1$ than in condition $2$. This situation violates the subset pivotality assumption, since the joint distribution of two or more features is changing across conditions while the marginal distributions of each of these features remains the same. In a situation like this it is possible that either $H_{1}$ or $H_{2}$ (or both) would be rejected more frequently than expected by chance using the max-T adjusted p-values.

There is no way to know if the subset pivotality assumption is accurate in our context, although we note that it is a strikingly singular manipulation to modify the joint distribution of a collection of variables with absolutely no modification of the marginal distributions of the variables. (In fact, it is impossible to do so if one also requires that the distributions of linear combinations of the variables remain unchanged.) For our scientific purposes, however, we think that the risk of violations of subset pivotality are small for the following reasons.

(1) The primary conclusion of our paper is that there are detectable electrophysiological differences between DBS used in different brain regions and/or using different stimulation parameters. This conclusion is unaffected by the subset pivotality assumption. Indeed, consider the global p-value

$$p_{*}=\frac{1}{L+1}\sum_{k=0}^{L} \mathbb{1}\left[ T_{*}^{k}\geq T_{*}^{0} \right]$$

This is simply a Monte Carlo permutation test p-value for comparing condition $1$ versus condition $2$ on the basis of the test statistic $T_{*}$, which uses all of the features. Since we have $p_{*}\leq p_{i}^{*}$ for all $i=1,\ldots,d$, then a rejection of any $H_{i}$ based on the adjusted p-value $p_{i}^{*}$ automatically implies a rejection of the null hypothesis that there is no difference between conditions.

(2) If the subset pivotality condition is not true, the only danger is that we will mistakenly conclude that differences in DBS create differences in some brain region (or frequency band) when, in fact, DBS only creates differences in that region’s (or band’s) relationship to another region (or band). It begins to border on philosophical whether this is even a mistaken scientific conclusion or not.

(3) Furthermore, since $p_{i}^{*}\geq p_{i}$ for each feature $i$, our adjusted p-values can only be used to reject null hypothesis $H_{i}$ when the data provide strong evidence against $H_{i}$ when viewed in isolation, i.e., when the $i$th feature measurements are noticeably different across conditions.

To test the effect of current-steering within each DBS lead on neural responses across the sampled networks, we performed a one-way ANOVA (using the *anova1.m* function in the statistics toolbox in MATLAB) for 30 categories comprised of each (region,band) combination using the five regions and six frequency bands of interest on the z-scored data in subject A. To demonstrate the effect of current steering on modulation of neural activity, we chose frequency bands with the greatest amount of variance across ROIs in response to current steered stimulation. We did this by first extracting the f-statistic values computed via a one-way ANOVA and then summed the f-statistic values for each frequency band across all ROIs and ranked the frequency bands in order of the highest f-statistic values. This resulted in alpha band as the highest ranked frequency band showing the greatest amount of variance across ROIs in response to current steering across the right SCC DBS lead, and theta band as the highest ranked frequency band showing the greatest amount of variance across ROIs in response to current steering across the right VC/VS DBS lead. Uncorrected p-values, f-statistic values and degrees of freedom are reported in Supplementary Table S7. While we performed a one-way anova to compute p-values to determine significance, due to the small number of trials per current configuration per DBS lead (N = 5) we did not perform additional statistical testing.

**Supplementary Tables**

| **Subject A** | | | | | | |
| --- | --- | --- | --- | --- | --- | --- |
| ROIs | Frequency | | | | | |
|  | Delta | Theta | Alpha | Beta | Low Gamma | High Gamma |
| dACC | 0.106 | 1.000 | 0.657 | 0.179 | 0.468 | 1.000 |
| Amygdala | 0.983 | 0.087 | 0.035 | 0.001 | 0.001 | 0.001 |
| mOFC | 0.123 | 0.182 | 0.001 | 0.019 | 0.229 | 0.190 |
| lOFC | 1.000 | 0.860 | 0.130 | 0.001 | 0.019 | 0.156 |
| vmPFC | 0.001 | 0.609 | **0.007** | 0.001 | 0.171 | 0.008 |
| **Subject B** | | | | | | |
| ROIs | Frequency | | | | | |
|  | Delta | Theta | Alpha | Beta | Low Gamma | High Gamma |
| dACC | 0.783 | 1.000 | 0.920 | 0.014 | 0.002 | 0.001 |
| Amygdala | 0.444 | 0.041 | 1.000 | 1.000 | 0.001 | 0.001 |
| mOFC | 0.663 | 0.147 | 0.787 | 0.999 | 0.132 | 0.032 |
| lOFC | 0.626 | 0.608 | 0.667 | 0.804 | 1.000 | 0.050 |
| vmPFC | 0.235 | 1.000 | 1.000 | 1.000 | 1.000 | 0.872 |

**Supplementary Table S1****:** Corrected p-values resulting from statistical testing between stimulation in the left SCC and baseline (pre-stim/baseline vs post-stimulation)

| **Subject A** | | | | | | |
| --- | --- | --- | --- | --- | --- | --- |
| ROIs | Frequency | | | | | |
|  | Delta | Theta | Alpha | Beta | Low Gamma | High Gamma |
| dACC | 0.021 | 1.000 | 1.000 | 0.553 | 0.999 | 1.000 |
| Amygdala | 1.000 | 0.763 | 0.001 | 0.001 | 0.001 | 0.001 |
| mOFC | 0.440 | 0.560 | 0.001 | 0.001 | 0.165 | 0.136 |
| lOFC | 0.997 | 0.964 | 0.710 | 0.001 | 0.001 | 0.007 |
| vmPFC | 0.001 | 0.266 | 0.038 | 0.001 | 1.000 | 0.314 |
| **Subject B** | | | | | | |
| ROIs | Frequency | | | | | |
|  | Delta | Theta | Alpha | Beta | Low Gamma | High Gamma |
| dACC | 0.511 | 1.000 | 1.000 | 0.003 | 0.108 | 0.001 |
| Amygdala | 0.538 | 0.001 | 0.160 | 0.005 | 0.001 | 0.001 |
| mOFC | 0.918 | 0.230 | 1.000 | 0.359 | 1.000 | 0.993 |
| lOFC | 0.918 | 0.572 | 1.000 | 0.368 | 1.000 | 0.751 |
| vmPFC | 0.068 | 0.951 | 0.822 | 0.292 | 1.000 | 1.000 |

**Supplementary Table S2:** Corrected p-values resulting from statistical testing between stimulation in the right SCC and baseline (pre-stim/baseline vs post-stimulation).

| **Subject A** | | | | | | |
| --- | --- | --- | --- | --- | --- | --- |
| ROIs | Frequency | | | | | |
|  | Delta | Theta | Alpha | Beta | Low Gamma | High Gamma |
| dACC | 0.020 | 0.945 | 1.000 | 1.000 | 0.917 | 1.000 |
| Amygdala | 0.969 | 0.308 | 0.948 | 0.593 | 0.063 | 0.018 |
| mOFC | 0.038 | 0.005 | 0.004 | 0.870 | 1.000 | 1.000 |
| lOFC | 0.565 | 0.079 | 0.100 | 1.000 | 0.152 | 0.994 |
| vmPFC | 0.002 | 0.070 | 1.000 | 1.000 | 0.824 | 0.762 |
| **Subject B** | | | | | | |
| ROIs | Frequency | | | | | |
|  | Delta | Theta | Alpha | Beta | Low Gamma | High Gamma |
| dACC | 0.927 | 0.944 | 0.291 | 0.001 | 0.019 | 0.001 |
| Amygdala | 0.215 | 0.062 | 1.000 | 0.027 | 0.592 | 0.084 |
| mOFC | 0.988 | 0.143 | 0.734 | 1.000 | 1.000 | 0.989 |
| lOFC | 1.000 | 0.870 | 0.999 | 0.995 | 1.000 | 1.000 |
| vmPFC | 0.123 | 1.000 | 0.984 | 1.000 | 0.400 | 0.253 |

**Supplementary Table S3:** Corrected p-values resulting from statistical testing between stimulation in the left VC/VS and baseline (pre-stim/baseline vs post-stimulation).

| **Subject A** | | | | | | |
| --- | --- | --- | --- | --- | --- | --- |
| ROIs | Frequency | | | | | |
|  | Delta | Theta | Alpha | Beta | Low Gamma | High Gamma |
| dACC | 0.856 | 0.023 | 1.000 | 0.965 | 1.000 | 1.000 |
| dlPFC | 1.000 | 0.991 | 0.908 | 0.991 | 0.028 | 0.017 |
| mOFC | 0.016 | 0.614 | 0.994 | 1.000 | 1.000 | 0.999 |
| lOFC | 1.000 | 1.000 | 1.000 | 0.994 | 0.999 | 0.748 |
| vmPFC | 0.004 | 0.023 | 1.000 | 0.863 | 0.215 | 0.961 |
| **Subject B** | | | | | | |
| ROIs | Frequency | | | | | |
|  | Delta | Theta | Alpha | Beta | Low Gamma | High Gamma |
| dACC | 1.000 | 0.884 | 0.662 | 0.003 | 1.000 | 0.001 |
| Amygdala | 0.473 | 0.438 | 1.000 | 0.092 | 0.999 | 0.001 |
| mOFC | 0.745 | 0.965 | 0.971 | 0.005 | 0.454 | 1.000 |
| lOFC | 0.380 | 0.666 | 1.000 | 0.002 | 0.253 | 1.000 |
| vmPFC | 0.855 | 0.997 | 1.000 | 0.001 | 0.029 | 0.575 |

**Supplementary Table S4:** Corrected p-values resulting from statistical testing between stimulation in the right VC/VS and baseline (pre-stim/baseline vs post-stimulation).

| **Subject A** | | | | | | |
| --- | --- | --- | --- | --- | --- | --- |
| ROIs | Frequency | | | | | |
|  | Delta | Theta | Alpha | Beta | Low Gamma | High Gamma |
| dACC | 0.983 | 1.000 | 0.529 | 0.216 | 1.000 | 0.988 |
| Amygdala | 1.000 | 1.000 | 0.489 | 0.058 | 0.024 | 0.046 |
| mOFC | 0.024 | 0.111 | 0.943 | 0.714 | 0.453 | 0.663 |
| lOFC | 0.355 | 0.987 | 1.000 | 0.003 | 0.001 | 0.009 |
| vmPFC | 0.030 | 1.000 | 0.026 | 0.001 | 0.001 | 0.001 |
| **Subject B** | | | | | | |
| ROIs | Frequency | | | | | |
|  | Delta | Theta | Alpha | Beta | Low Gamma | High Gamma |
| dACC | 1.000 | 0.921 | 0.984 | 0.422 | 0.877 | 0.903 |
| Amygdala | 1.000 | 1.000 | 1.000 | 0.038 | 0.001 | 0.001 |
| mOFC | 0.994 | 1.000 | 1.000 | 0.970 | 0.020 | 0.001 |
| lOFC | 0.981 | 1.000 | 0.994 | 1.000 | 1.000 | 0.003 |
| vmPFC | 1.000 | 1.000 | 0.982 | 1.000 | 0.075 | 0.001 |

**Supplementary Table S5:** Corrected p-values resulting from statistical testing between stimulation in the left SCC and stimulation in the left VC/VS (baseline-subtracted post-stimulation windows used for both DBS leads)

| **Subject A** | | | | | | |
| --- | --- | --- | --- | --- | --- | --- |
| ROIs | Frequency | | | | | |
|  | Delta | Theta | Alpha | Beta | Low Gamma | High Gamma |
| dACC | 1.000 | 0.137 | 1.000 | 0.009 | 1.000 | 1.000 |
| Amygdala | 1.000 | 1.000 | 0.620 | 0.002 | 0.002 | 0.001 |
| mOFC | 0.300 | 1.000 | 0.414 | 0.004 | 0.058 | 0.837 |
| lOFC | 0.990 | 0.991 | 0.992 | 0.047 | 0.004 | 0.391 |
| vmPFC | 0.798 | 0.998 | 0.071 | 0.001 | 0.012 | 0.004 |
| **Subject B** | | | | | | |
| ROIs | Frequency | | | | | |
|  | Delta | Theta | Alpha | Beta | Low Gamma | High Gamma |
| dACC | 0.001 | 0.694 | 0.543 | 1.000 | 0.015 | 1.000 |
| Amygdala | 1.000 | 0.010 | 0.264 | 1.000 | 0.001 | 1.000 |
| mOFC | 0.003 | 0.725 | 0.768 | 0.549 | 0.024 | 0.491 |
| lOFC | 0.002 | 1.000 | 1.000 | 0.209 | 0.336 | 0.217 |
| vmPFC | 0.012 | 1.000 | 0.998 | 0.097 | 0.025 | 0.278 |

**Supplementary Table S6:** Corrected p-values resulting from statistical testing between stimulation in the right SCC and stimulation in the right VC/VS (baseline-subtracted post-stimulation windows used for both DBS leads)

| **Alpha power following stimulation** | | | | **Theta power following stimulation** | | | |
| --- | --- | --- | --- | --- | --- | --- | --- |
| **R-SCC** | | | | **R-SCC** | | | |
| ROI | df | f-statistic | p-value | ROI | df | f-statistic | p-value |
| dACC | 6 | 0.54381508 | 0.770375 | dACC | 4 | 1.288509 | 0.294536 |
| Amygdala | 6 | 1.18697813 | 0.341678 | Amygdala | 4 | 1.402519 | 0.248526 |
| mOFC | 6 | 4.41423266 | 0.002922 | mOFC | 4 | 2.071399 | 0.088964 |
| lOFC | 6 | 3.48675677 | 0.010604 | lOFC | 4 | 1.278878 | 0.29875 |
| vmPFC | 6 | 1.24121801 | 0.315743 | vmPFC | 4 | 1.763105 | 0.143325 |
| **R-VC/VS** | | | | **R-VC/VS** | | | |
| ROI | df | f-statistic | p-value | ROI | df | f-statistic | p-value |
| dACC | 4 | 1.33490557 | 0.291522 | dACC | 4 | 2.570807 | 0.069391 |
| Amygdala | 4 | 0.2065304 | 0.931776 | Amygdala | 4 | 1.161635 | 0.357213 |
| mOFC | 4 | 0.43197588 | 0.783873 | mOFC | 4 | 1.003923 | 0.428749 |
| lOFC | 4 | 0.41683244 | 0.794511 | lOFC | 4 | 3.34029 | 0.030012 |
| vmPFC | 4 | 0.57318741 | 0.685224 | vmPFC | 4 | 0.634125 | 0.644018 |

**Supplementary Table S7:** Results from one-way ANOVA statistical testing of different current steering configurations across the right SCC and VC/VS DBS leads on alpha and theta power in ROIs (performed on z-scored data) in subject A. P-values are uncorrected.

**Supplementary Tables with uncorrected p-values following permutation testing**

| **Subject A** | | | | | | |
| --- | --- | --- | --- | --- | --- | --- |
| ROIs | Frequency | | | | | |
|  | Delta | Theta | Alpha | Beta | Low Gamma | High Gamma |
| dACC | 0.006 | 0.660 | 0.052 | 0.006 | 0.029 | 0.361 |
| Amygdala | 0.193 | 0.006 | 0.001 | 0.001 | 0.001 | 0.001 |
| mOFC | 0.006 | 0.012 | 0.001 | 0.002 | 0.015 | 0.009 |
| lOFC | 0.518 | 0.097 | 0.006 | 0.001 | 0.002 | 0.006 |
| vmPFC | 0.001 | 0.048 | 0.002 | 0.001 | 0.008 | 0.001 |
| **Subject B** | | | | | | |
| ROIs | Frequency | | | | | |
|  | Delta | Theta | Alpha | Beta | Low Gamma | High Gamma |
| dACC | 0.073 | 0.994 | 0.158 | 0.002 | 0.001 | 0.001 |
| Amygdala | 0.036 | 0.002 | 0.730 | 0.461 | 0.001 | 0.001 |
| mOFC | 0.054 | 0.007 | 0.096 | 0.378 | 0.011 | 0.004 |
| lOFC | 0.048 | 0.050 | 0.057 | 0.090 | 0.887 | 0.001 |
| vmPFC | 0.009 | 0.733 | 0.846 | 0.600 | 0.819 | 0.119 |

**Supplementary Table S8:** Uncorrected p-values resulting from statistical testing between stimulation in the left SCC and baseline (pre-stim/baseline vs post-stimulation)

| **Subject A** | | | | | | |
| --- | --- | --- | --- | --- | --- | --- |
| ROIs | Frequency | | | | | |
|  | Delta | Theta | Alpha | Beta | Low Gamma | High Gamma |
| dACC | 0.002 | 0.742 | 0.561 | 0.036 | 0.279 | 0.350 |
| Amygdala | 0.342 | 0.072 | 0.001 | 0.001 | 0.001 | 0.001 |
| mOFC | 0.016 | 0.050 | 0.001 | 0.001 | 0.011 | 0.007 |
| lOFC | 0.232 | 0.146 | 0.057 | 0.001 | 0.001 | 0.001 |
| vmPFC | 0.001 | 0.029 | 0.004 | 0.001 | 0.399 | 0.021 |
| **Subject B** | | | | | | |
| ROIs | Frequency | | | | | |
|  | Delta | Theta | Alpha | Beta | Low Gamma | High Gamma |
| dACC | 0.023 | 0.906 | 0.872 | 0.001 | 0.005 | 0.001 |
| Amygdala | 0.031 | 0.001 | 0.007 | 0.002 | 0.001 | 0.001 |
| mOFC | 0.150 | 0.006 | 0.606 | 0.024 | 0.670 | 0.292 |
| lOFC | 0.146 | 0.044 | 0.705 | 0.041 | 0.651 | 0.066 |
| vmPFC | 0.001 | 0.175 | 0.107 | 0.019 | 0.640 | 0.828 |

**Supplementary Table S9:** Uncorrected p-values resulting from statistical testing between stimulation in the right SCC and baseline (pre-stim/baseline vs post-stimulation).

| **Subject A** | | | | | | |
| --- | --- | --- | --- | --- | --- | --- |
| ROIs | Frequency | | | | | |
|  | Delta | Theta | Alpha | Beta | Low Gamma | High Gamma |
| dACC | 0.003 | 0.143 | 0.876 | 0.731 | 0.115 | 0.696 |
| Amygdala | 0.174 | 0.016 | 0.139 | 0.035 | 0.004 | 0.001 |
| mOFC | 0.001 | 0.001 | 0.001 | 0.092 | 0.743 | 0.511 |
| lOFC | 0.034 | 0.001 | 0.005 | 0.692 | 0.014 | 0.242 |
| vmPFC | 0.001 | 0.004 | 0.480 | 0.555 | 0.091 | 0.060 |
| **Subject B** | | | | | | |
| ROIs | Frequency | | | | | |
|  | Delta | Theta | Alpha | Beta | Low Gamma | High Gamma |
| dACC | 0.138 | 0.173 | 0.017 | 0.001 | 0.001 | 0.001 |
| Amygdala | 0.007 | 0.005 | 0.476 | 0.001 | 0.035 | 0.004 |
| mOFC | 0.264 | 0.007 | 0.066 | 0.712 | 0.920 | 0.287 |
| lOFC | 0.390 | 0.113 | 0.368 | 0.329 | 0.622 | 0.903 |
| vmPFC | 0.003 | 0.561 | 0.258 | 0.828 | 0.036 | 0.015 |

**Supplementary Table S10:** Uncorrected p-values resulting from statistical testing between stimulation in the left VC/VS and baseline (pre-stim/baseline vs post-stimulation).

| **Subject A** | | | | | | |
| --- | --- | --- | --- | --- | --- | --- |
| ROIs | Frequency | | | | | |
|  | Delta | Theta | Alpha | Beta | Low Gamma | High Gamma |
| dACC | 0.096 | 0.002 | 0.453 | 0.134 | 0.707 | 0.491 |
| dlPFC | 0.916 | 0.183 | 0.132 | 0.190 | 0.002 | 0.003 |
| mOFC | 0.001 | 0.039 | 0.212 | 0.835 | 0.507 | 0.251 |
| lOFC | 0.582 | 0.737 | 0.358 | 0.222 | 0.229 | 0.063 |
| vmPFC | 0.001 | 0.002 | 0.861 | 0.084 | 0.012 | 0.132 |
| **Subject B** | | | | | | |
| ROIs | Frequency | | | | | |
|  | Delta | Theta | Alpha | Beta | Low Gamma | High Gamma |
| dACC | 0.822 | 0.108 | 0.054 | 0.001 | 0.830 | 0.001 |
| Amygdala | 0.018 | 0.026 | 0.689 | 0.007 | 0.369 | 0.001 |
| mOFC | 0.075 | 0.205 | 0.231 | 0.001 | 0.029 | 0.524 |
| lOFC | 0.020 | 0.055 | 0.465 | 0.001 | 0.012 | 0.720 |
| vmPFC | 0.085 | 0.354 | 0.460 | 0.001 | 0.002 | 0.052 |

**Supplementary Table S11:** Uncorrected p-values resulting from statistical testing between stimulation in the right VC/VS and baseline (pre-stim/baseline vs post-stimulation).

| **Subject A** | | | | | | |
| --- | --- | --- | --- | --- | --- | --- |
| ROIs | Frequency | | | | | |
|  | Delta | Theta | Alpha | Beta | Low Gamma | High Gamma |
| dACC | 0.180 | 0.251 | 0.041 | 0.012 | 0.732 | 0.196 |
| Amygdala | 0.948 | 0.719 | 0.021 | 0.004 | 0.002 | 0.003 |
| mOFC | 0.002 | 0.006 | 0.144 | 0.069 | 0.030 | 0.066 |
| lOFC | 0.019 | 0.208 | 0.955 | 0.001 | 0.001 | 0.001 |
| vmPFC | 0.004 | 0.432 | 0.001 | 0.001 | 0.001 | 0.001 |
| **Subject B** | | | | | | |
| ROIs | Frequency | | | | | |
|  | Delta | Theta | Alpha | Beta | Low Gamma | High Gamma |
| dACC | 0.287 | 0.114 | 0.195 | 0.022 | 0.092 | 0.082 |
| Amygdala | 0.413 | 0.354 | 0.679 | 0.002 | 0.001 | 0.001 |
| mOFC | 0.221 | 0.822 | 0.806 | 0.156 | 0.003 | 0.001 |
| lOFC | 0.193 | 0.563 | 0.221 | 0.374 | 0.325 | 0.001 |
| vmPFC | 0.494 | 0.718 | 0.189 | 0.417 | 0.002 | 0.001 |

**Supplementary Table S12:** Uncorrected p-values resulting from statistical testing between stimulation in the left SCC and stimulation in the left VC/VS (baseline-subtracted post-stimulation windows used for both DBS leads)

| **Subject A** | | | | | | |
| --- | --- | --- | --- | --- | --- | --- |
| ROIs | Frequency | | | | | |
|  | Delta | Theta | Alpha | Beta | Low Gamma | High Gamma |
| dACC | 0.471 | 0.005 | 0.850 | 0.002 | 0.612 | 0.828 |
| Amygdala | 0.367 | 0.426 | 0.056 | 0.001 | 0.001 | 0.001 |
| mOFC | 0.021 | 0.511 | 0.032 | 0.001 | 0.004 | 0.084 |
| lOFC | 0.203 | 0.203 | 0.215 | 0.003 | 0.001 | 0.029 |
| vmPFC | 0.059 | 0.261 | 0.002 | 0.001 | 0.001 | 0.001 |
| **Subject B** | | | | | | |
| ROIs | Frequency | | | | | |
|  | Delta | Theta | Alpha | Beta | Low Gamma | High Gamma |
| dACC | 0.001 | 0.042 | 0.029 | 0.800 | 0.002 | 0.345 |
| Amygdala | 0.934 | 0.002 | 0.017 | 0.379 | 0.001 | 0.729 |
| mOFC | 0.001 | 0.059 | 0.081 | 0.039 | 0.002 | 0.027 |
| lOFC | 0.001 | 0.625 | 0.619 | 0.012 | 0.025 | 0.011 |
| vmPFC | 0.001 | 0.593 | 0.258 | 0.003 | 0.002 | 0.015 |

**Supplementary Table S13:** Uncorrected p-values resulting from statistical testing between stimulation in the right SCC and stimulation in the right VC/VS (baseline-subtracted post-stimulation windows used for both DBS leads)
